# 5-HT_2C_ receptors in the nucleus accumbens constrain the rewarding effects of MDMA

**DOI:** 10.1101/2024.10.20.619256

**Authors:** Matthew B. Pomrenze, Sam Vaillancourt, Juliana S. Salgado, Kendall B. Raymond, Pierre Llorach, Gavin C. Touponse, Daniel F. Cardozo Pinto, Zahra Rastegar, Austen B. Casey, Neir Eshel, Robert C. Malenka, Boris D. Heifets

## Abstract

MDMA is a promising adjunct to psychotherapy and has well-known abuse liability, although less than other amphetamine analogs. While the reinforcing dopamine (DA)-releasing properties of MDMA are on par with methamphetamine (METH), MDMA is a far more potent serotonin (5-HT) releaser, via the 5-HT transporter (SERT). MDMA-mediated 5-HT release in a major reward center, the nucleus accumbens (NAc), drives prosocial behaviors via 5-HT1BR activation. We hypothesized that this prosocial mechanism contributes to the reduced reinforcing properties of MDMA compared to METH and used a platform of assays to predict the balance of prosocial and abuse-linked effects of (R)-MDMA, a novel entactogen in clinical development. NAc DA release, measured by GRAB-DA photometry in vivo, increased in proportion to MDMA (7.5 and 15 mg/kg, i.p.) and METH (2 mg/kg i.p.)-conditioned place preference (CPP). Using conditional knockouts (cKOs) for DAT and SERT, microdialysis, and photometry, we found that MDMA-released 5-HT limited MDMA-released DA through actions in the NAc, rather than at ventral tegmental area DAergic cell bodies. SERT cKO reduced the MDMA dose required for CPP three-fold. This enhanced MDMA-CPP and increased DA release were replicated by intra-NAc infusion of either a 5-HT reuptake inhibitor (escitalopram) to prevent MDMA interaction with SERT, or a 5-HT2CR antagonist (SB242084), but not by the 5-HT1BR antagonist NAS-181. These data support separate mechanisms for the low abuse potential versus prosocial effect of MDMA. Using this platform of assays, (R)-MDMA is predicted to have prosocial effects and low abuse potential.

## INTRODUCTION

MDMA (also known as ‘molly’ or ‘ecstasy’) shows promise as an adjunct to psychotherapy for posttraumatic stress disorder (PTSD) [1, 2]. The efficacy of MDMA assisted therapy may stem from MDMA’s unique behavioral properties, which include an enhanced sense of emotional connectedness and empathy, and reduced fear when confronted with aversive stimuli, including traumatic memories [3]. MDMA, an amphetamine analog, is also prone to misuse and abuse [4], an important risk consideration for treating patients with PTSD, many of whom have comorbid substance use disorders [5]. However, MDMA is not as widely abused as closely related drugs in the amphetamine class, such as methamphetamine (METH; [6, 7]), raising the possibility that MDMA’s reduced abuse potential is mechanistically linked to its therapeutic behavioral effects.

The neurochemical mechanism of MDMA differs from that of METH in at least one major aspect: while METH, MDMA and other amphetamines share an affinity for the dopamine transporter (DAT), driving non-vesicular DA release by reverse DA transport [8], MDMA also has a uniquely high affinity for the 5-HT transporter (SERT), driving supraphysiological 5-HT release via reverse transport [9]. We and others have found that in a major brain reward center, the nucleus accumbens (NAc), 5-HT release via an MDMA / SERT interaction and subsequent activation of the 5-HT_1B_ receptor (5-HT_1B_R) accounts for MDMA’s prosocial effects in a variety of social behavioral assays [10–14]. These prosocial behaviors, including social approach in the three-chamber test (3-CT), social transfer of affective states, and social reward learning all reflect behavioral processes modified by MDMA in human studies [15–17]. In contrast, nonsocial drug reward evoked by METH and higher doses of MDMA has consistently been attributed to DA release in the same brain structure, the NAc, across species [10, 18].

The apparent relationship between SERT affinity, 5-HT release, and reduced abuse liability among amphetamine analogs and other stimulants has long been appreciated [7, 19, 20]. Investigations focused on brain-region specific regulation of DA release suggested a variety of candidate processes, including suppression of glutamatergic cortical inputs to dorsal striatum via 5-HT_1B_R [21], modulation of dopaminergic cell body excitability in the ventral tegmental area (VTA) via 5-HT_1B_R [22, 23] and 5-HT_2C_R [24–26], and modulation of DA release in the NAc by 5-HT_1B_R [27, 28], 5-HT_2A_R [29], 5-HT_2B_R [30, 31] and 5-HT_2C_Rs [29, 32]. Conflicting results with some 5-HTR ligands have added to the uncertainty about how stimulant-evoked 5-HT and DA release interact [29, 33].

Building on our prior work showing that prosocial and abuse-linked properties of MDMA are mediated by 5-HT and DA release in the NAc, respectively, we now focus on the relationship between 5-HT and DA released by MDMA, and the specific brain regions and 5-HTRs that mediate their interaction. The goals of this study were threefold: 1) establish whether 5-HT release wholly accounts for the difference in abuse-related properties of racemic MDMA and METH; 2) isolate the site and receptor-specific action of 5-HT on DA release at a selectively prosocial MDMA dose; and 3) test predictions about the comparative prosocial versus rewarding effects of (*R*)-MDMA, which is being actively developed for clinical use as an entactogen[34], a drug class that induces feelings of empathy and emotional openness. We use fluorescent Ca^2+^ and neuromodulator sensors *in vivo*, brain-region specific drug infusions, DAT and SERT conditional knockout mice (cKO), and a simple, widely-used assay, conditioned place preference (CPP) to establish that MDMA-induced activation of 5-HT_2C_Rs in the NAc, but not 5-HT_1B_Rs, actively suppresses DA release in a manner that is strongly associated with the reward-related properties of MDMA. Having found strong relationships between NAc DA release and the ability to induce CPP, and NAc 5-HT release and increased social approach in the 3-CT, we use these simple *in vivo* physiological assays with (*R*)-MDMA. We accurately predicted that (*R*)-MDMA has prosocial properties and limited abuse liability in mice, suggesting these assays’ utility in identifying novel low-risk entactogens.

## MATERIALS AND METHODS

### Subjects

Male and female C57BL/6J (Jackson Laboratory, stock #00664), aged 8 to 16 weeks old and the following transgenic lines were used:

1. Heterozygous *7630403G23Rik^Tg(Th-cre)1Tmd^*/J (TH-Cre) mice (Jackson Laboratory, stock #008601).
2. Heterozygous *Slc32a1^tm2(cre)Lowl^*/MwarJ (*Vgat*-Cre) mice (Jackson Laboratory, stock #028862)
3. Conditional DAT KO (floxed *Slc6a3* [DAT^fl/fl^]; DAT cKO) mice were derived from *Slc6a3^tm1a(KOMP)Wtsi^*mice (UC Davis KOMP repository line, RRID:MMRRC_062518-UCD), by crossing to a FLPo deleter strain (Jackson Laboratory, stock #012930), and back crossing to wild type C57BL/6J to remove the FLP gene. Inducible homozygous variants of the DAT cKO mouse were generated by crossing DAT cKO mice with *Th^tm1.1(cre/ERT2)Ddg^*/J (TH-Cre^ERT2^) mice (Jackson Laboratory, stock #025614).
4. Conditional SERT KO (floxed *Slc6a4* [SERT^fl/fl^]; SERT cKO) mice were a gift from J.-Y. Sze, Albert Einstein College of Medicine, [35]. Inducible variants of the SERT cKO mouse were generated by crossing SERT^fl/fl^ mice with C57BL/6-Tg(Nes-cre/ERT2)KEisc/J (Nestin-Cre^ERT2^) mice (Jackson Laboratory, stock #016261).

All mice were kept on a C57BL/6J background and group housed on a 12-hr light/dark cycle with food and water *ad libitum*. All procedures complied with animal care standards set forth by the National Institute of Health and were approved by Stanford University’s Administrative Panel on Laboratory Animal Care and Administrative Panel of Biosafety.

### Viral vectors

AAV9-hSyn-GRAB-DA4.4 (DA2m) and AAV9-hSyn-GRAB-5HT3.6 (5HT3) were purchased from WZ Biosciences (Columbia, MD). AAV-hSyn-FLEX-GCaMP8m was purchased from Addgene. All viruses were injected at 4-6 · 10^12^ infectious units per mL.

### Stereotaxic surgery

#### Virus injection and optical fiber implants

Mice of at least 8 weeks of age were anesthetized with isoflurane (1-2% v/v) and secured in a stereotaxic frame (David Kopf Instruments, Tujunga, CA). Viruses were injected unilaterally into the NAc medial shell (AP +1.2, ML +0.7, DV −3.6 from brain surface) or VTA (AP −3.3, ML +0.3, DV −4.1 from skull) at a rate of 150 nL min^-1^ (800 nL total volume) with a borosilicate pipette coupled to a pump-mounted 5 µL Hamilton syringe. Injector pipettes were slowly retracted after a 5 min diffusion period. Optical fibers (Doric Lenses) with a 400 μm core and 0.66 NA were unilaterally implanted over the NAc (AP +1.2, ML +0.7, DV −3.5 from brain surface) or VTA (AP −3.3, ML +0.3, DV −4.0 from skull). Optical fibers were secured to the skull with stainless steel screws (thread size 00-90 x 1/16, Antrin Miniature Specialties), C&B Metabond, and light-cured dental adhesive cement (Geristore A&B paste, DenMat). Mice were group housed to recover for at least 3 weeks before recordings began.

#### Cannula implants

For drug microinfusions, a 26-gauge threaded bilateral guide cannula (P1 Technologies), 3.5 mm from the cannula base, was implanted over the NAc (AP +1.2; ML ±0.7; DV −3.1 from brain surface). For microdialysis, a bilateral guide cannula (Amuza, CXG-04) was implanted over the NAc (AP +1.2; ML ±0.7; DV −3.1). For drug microinfusions before photometry recordings, a dual optical fiber-cannula (Doric Lenses, OmFC, fiber with 400 um core and 0.66 NA, 25-gauge cannula) was implanted over the NAc (AP +1.2, ML +0.7, DV −3.5 from brain surface). Implants were secured to the skull with stainless steel screws (thread size 00-90 x 1/16, Antrin Miniature Specialties), C&B Metabond, and light-cured dental adhesive cement (Geristore A&B paste, DenMat). Mice were group housed to recover for at least 3 weeks before experiments began.

### Microdialysis

A bilateral microdialysis probe (CX-1-04-01, Amuza,CA) was inserted into a cannula implanted 48 hr previously, over the NAc. Mice were then placed in a clean arena and connected to a swivel arm (FC-90, Amuza,CA) coupled to the sample collector. Artificial cerebrospinal fluid (aCSF) was injected at a rate of 1 μl/min (Microdialysis pump ESP-180LD, Amuza, CA). Baselines were collected for 40 minutes and then dialysate was collected for 100 min after an injection of MDMA (7.5 mg/kg, ip). Dialysates were analyzed by HPLC at the end of each session.

### Drug administration

(±)-MDMA (5, 7.5, or 15 mg/kg, Organix or NIDA Drug Supply Program), (*R*)-MDMA (10, 20, or 40 mg/kg, NIDA Drug Supply Program), methamphetamine (METH, 2 mg/kg, Sigma-Aldrich), cocaine (15 mg/kg, Sigma-Aldrich), and D-fenfluramine (FEN, 10 mg/kg, Tocris) were dissolved in saline and administered intraperitoneally (ip) at a volume of 10 mL/kg. For intra-NAc infusions, escitalopram oxalate (Scit, 0.5 μg in 0.5 μL, Tocris) and NAS-181 (0.5 μg in 0.5 μL, Tocris) were dissolved in saline. The 5-HT_2C_ receptor antagonist SB242084 (1 μM in 0.5 μL, Tocris) was dissolved in DMSO and diluted in saline (0.01% v/v DMSO). All drugs were microinjected into the NAc 10 min before an injection of MDMA (5 mg/kg, ip) or saline. Tamoxifen (T5648, Sigma-Aldrich) was dissolved in corn oil, vortexed on heat, and injected (75 mg/kg, ip) for three consecutive days. Experiments were performed at least one week later.

For intracranial infusions before behavior, the bilateral cannula stylet was removed, and a 33-gauge bilateral injector was inserted into the cannula. Drugs were microinjected to a total volume of 500 nL at a rate of 150 nL/min. Injectors were left in place for 2 min following microinjections. Mice were then injected with MDMA and conditioned in CPP chambers. For infusions before photometry recordings, the unilateral cannula stylet was removed from the optical fiber-cannula implant and a 100 um core polyimide injector (FI_OmFC-ZF_100/170) was inserted through the cannula. Drugs were microinjected to a total volume of 500 nL at a rate of 150 nL/min. Injectors were left in place for 2 min following microinjections. Mice were then immediately transferred to the recording room, connected to optical patch cords, and the recording began.

### Conditioned place preference (CPP)

To evaluate drug reinforcing effects, mice were allowed to explore a 2-sided CPP chamber with distinct tactile floors and wall patterns (Med Associates Inc.) in a 15 min pretest. The next day, mice were confined to one side for 30 min after receiving an injection of saline. The next day, mice were confined to the opposite side of the chamber immediately after receiving an injection of MDMA (5, 7.5, or 15 mg/kg, ip), (*R*)-MDMA (20 mg/kg, ip), METH (2 mg/kg, ip), FEN (10 mg/kg, ip), or cocaine (15 mg/kg, ip) for 30 min. This was repeated once again for a total of two drug conditioning sessions. 24 hr after the last conditioning session, mice were allowed to explore both sides of the CPP chamber during a 15 min posttest. Preference was calculated as the percentage of time spent in the drug-paired side of the chamber during the posttest. Drug-paired sides were assigned in a counterbalanced and unbiased fashion such that the average preference for the drug-paired side during the pretest was ∼50% for all groups.

### 3-chamber sociability test (3-CT)

3-chamber sociability testing was performed in an arena with three separate chambers as previously described [10]. On day one, test mice were habituated to the arena containing two empty wire mesh cups placed in the two outer chambers for 5 min. Conspecific mice were also habituated to the mesh cups for 5 min. On day two, a conspecific mouse (age, strain- and sex-matched) was placed into one of the wire mesh cups and test mice were injected with MDMA (7.5 mg/kg, ip), (*R*)-MDMA (20 mg/kg, ip), FEN (10 mg/kg, ip), or METH (2 mg/kg, ip) and placed into the center chamber. After a 15 min habituation, the barriers were raised, and the test mouse was allowed to explore freely during a 30 min session. The placement of the conspecific mouse was counterbalanced across sessions. Location of mice was assayed automatically using video tracking software (BIOBSERVE). Sociability was calculated as: ((time in social side – time in empty side) / (time in social side + time in empty side))*100.

### Fiber photometry

Fiber photometry was performed as previously described [10, 12, 36, 37]. AAV9-hSyn-GRAB_DA_, AAV9-hSyn-GRAB_5-HT_, or AAV9-hSyn-FLEX-GCaMP8m were injected into the NAc or VTA with a fiber directed above. After at least 3 weeks, mice were habituated to the photometry setup. On the test day, mice were connected to patch cables and allowed to habituate alone in the homecage for 10 min. Mice were then injected with the appropriate drug and recordings continued for another 40 min. As outlined above, mice that received microinjections did so in a separate room immediately before being subjected to the recording procedure.

Data were acquired using Synapse software controlling an RZ5P lock-in amplifier (Tucker-Davis Technologies). GRAB and GCaMP8m sensors were excited by frequency-modulated 465 and 405 nm LEDs (Doric Lenses). Optical signals were band-pass-filtered with a fluorescence mini cube (Doric Lenses) and signals were digitized at 6 kHz. Signal processing was performed with custom scripts in MATLAB (MathWorks) [10, 12, 36, 37]. Briefly, for GRAB_DA_ and GRAB_5-HT_ experiments, 100Hz down-sampled signals were de-bleached by fitting a mono-exponential decay function to the baseline (pre-drug) portion of the signal and subtracting this fit curve from the full-length trace. To calculate ΔF/F, all fluorescence intensity values (F_465_) for the entire time course were referenced to the mean (F_mean_) fluorescence for all values of the de-bleached pre-drug baseline as (F_465_ − F_mean_) / F_mean_. Z-scoring was performed similarly, using only the pre-drug baseline to calculate F_mean_ and F_stdev_ (Z = [F_465_ – F_mean_] / F_stdev_). For GCaMP8m experiments, the entire time course was used for curve-fitting and calculating F_mean_ and F_stdev_. The resulting traces were smoothed (MATLAB, ‘filtfilt’) using a zero-phase moving average filter with an equally weighted 100 sample window. Transient detection was automated (MATLAB Signal Processing Toolbox, ‘findpeaks’), with a 3 sec lockout between events. Transients were defined as Z-scored fluorescence from −3 sec to +4 sec relative to the detected peak. Each transient was normalized to its baseline defined as −2.9 to −2 sec. Area under the curve was defined as the integral between 0 and 40 min. All photometry data was processed blind to condition with the same method.

### Immunohistochemistry

Mice were transcardially perfused with 4% paraformaldehyde in PBS, and brains were postfixed overnight in the same solution. Coronal brain sections (40 μM) were cut on a vibratome and stored in cold PBS. Free-floating sections were washed three times in PBS containing 0.2% Triton X-100 (PBST) for 10 min before incubation in a blocking solution containing PBST and 3% normal goat serum (NGS) for 1 hr, rocking at room temperature. Sections were then incubated in blocking solution with primary antibodies rocking at 4°C for 20 hrs. Primary antibodies used were mouse anti-TH (1:1000, Immunostar, 22941) and chicken anti-GFP (1:1000, Aves Labs, GFP-1010). Sections were then washed in PBST three times for 10 min and then incubated in the same blocking solution containing species-specific secondary antibodies (1:700, Alexa Fluor 647 goat anti-mouse and Alexa Fluor 488 goat anti-chicken). Sections were then rinsed three times in PBS for 5 min and mounted onto Superfrost slides (Fisher Scientific) with Fluoromount-G containing DAPI (SouthernBiotech). Slides were kept in the dark at 4°C until imaging on a Nikon A1 confocal microscope.

### Statistical analysis

Investigators were blinded to the manipulations experimental subjects had received for behavioral assays and photometry recordings. All behavioral data were analyzed and graphed with GraphPad Prism 9. All photometry data were processed and analyzed in MATLAB with custom scripts. Data distribution and variance were tested using Shapiro-Wilk normality tests. Normally distributed data were analyzed by unpaired, two-tailed t-tests, or one- or two-factor ANOVA with *post-hoc* Sidak, Tukey, or Dunnett correction for multiple comparisons. Paired comparisons were performed when appropriate (e.g. before versus after conditioning). Differences were considered significant when *P* < 0.05. All pooled data are expressed as mean ± SEM.

## RESULTS

### Dissociation of MDMA effects on sociability and reward via DA and 5-HT release

To quantify 5-HT and DA release in the NAc associated with MDMA-evoked prosocial behavior and drug reward, we first reproduced our and others’ work demonstrating behaviorally selective doses of MDMA (7.5 mg/kg, ip) that was previously shown to be prosocial but not reinforcing in wild-type mice [10, 11, 13, 14, 38, 39] and compared its effects on social behavior and reward to those triggered by d-Fenfluramine (FEN, 10 mg/kg, ip) and methamphetamine (METH, 2 mg/kg, ip), two drugs with preferential actions at SERT and DAT respectively [19]. In the 3-CT, FEN and MDMA (7.5 mg/kg) significantly increased sociability to a similar extent, however METH was ineffective (Fig. 1a-c) [10]. In contrast to these effects on social behavior, in a conditioned place preference (CPP) assay, mice failed to show a preference for contexts paired with MDMA (7.5 mg/kg) or FEN, but did so with METH and a higher dose of MDMA (15 mg/kg; Fig. 1d-f), indicative of a reinforcing drug effect.

**Figure 1.**
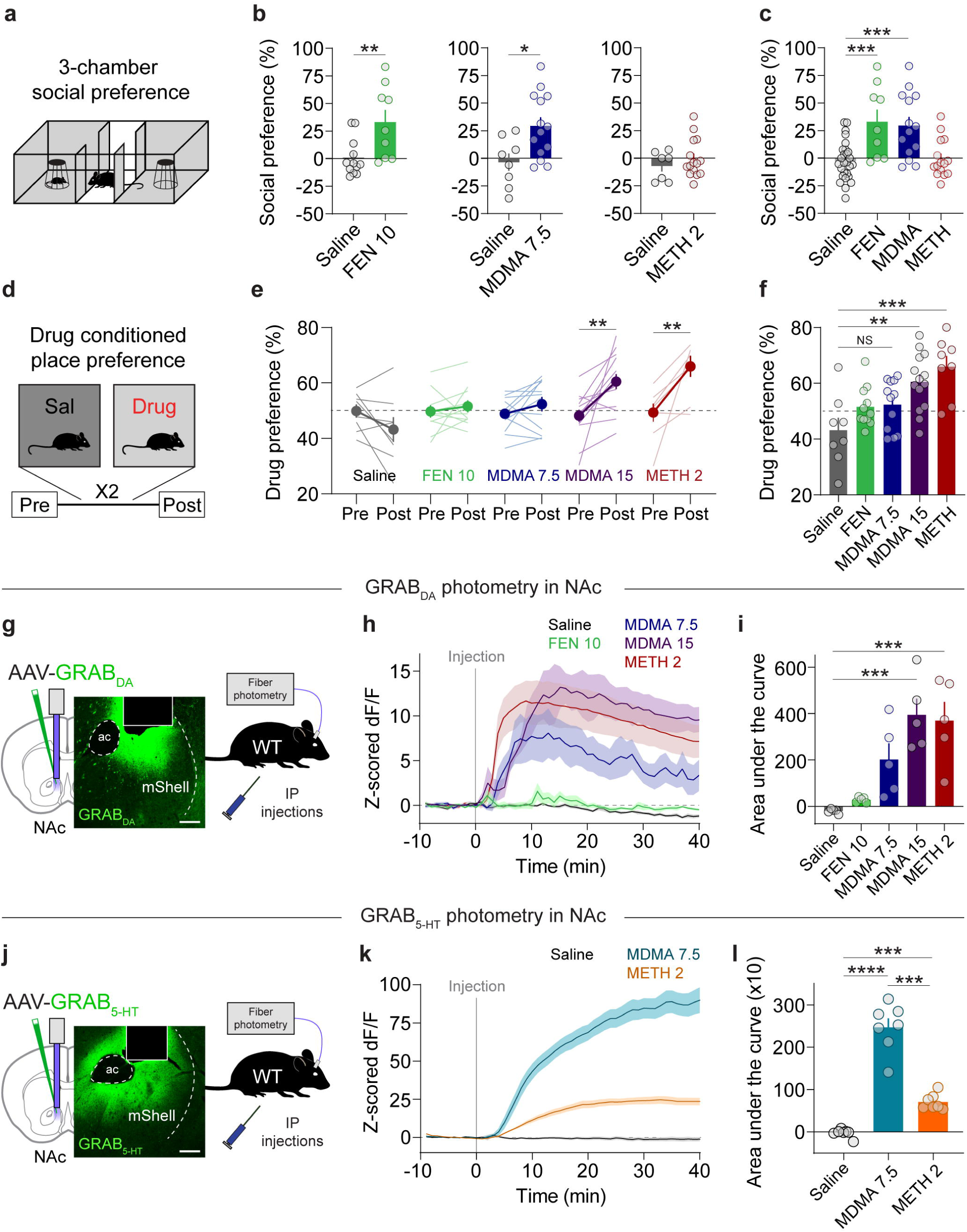
MDMA engages distinct reward processes. **a.** Schematic of 3-chamber social preference test. **b.** Left, effect of FEN (10 mg/kg) on social preference in the 3-chamber test. Unpaired, two-tailed t-test, t_19_ = 2.946, **p = 0.0083. Center, effect of MDMA (7.5 mg/kg) on social preference. Unpaired, two-tailed t-test, t_20_ = 2.746, *p = 0.0125. Right, effect of METH (2 mg/kg) on social preference. Unpaired, two-tailed t-test, t_20_ = 0.921, p = 0.3681. **c.** Summary of social preference across different drugs. Ordinary one-way ANOVA, F_3,61_ = 10.11, ****p < 0.0001. Dunnett’s multiple comparisons test, saline vs FEN ***p = 0.0004, saline vs MDMA ***p = 0.0002, saline vs METH p = 0.9831. **d.** Schematic of drug conditioned place preference (CPP) testing and protocol. **e.** Preference for each drug-paired context for each drug. Saline: paired, two-tailed t-test, t_7_ = 1.799, p = 0.115. FEN: paired, two-tailed t-test, t_10_ = 0.739, p = 0.4768. MDMA 7.5: paired, two-tailed t-test, t_11_ = 1.459, p = 0.1726. MDMA 15: paired, two-tailed t-test, t_13_ = 3.672, **p = 0.0028. METH 2: paired, two-tailed t-test, t_7_ = 3.714, **p = 0.0075. **f.** Summary and direct comparison of CPP scores between drug groups. Ordinary one-way ANOVA, F_4,48_ = 6.965, ***p = 0.0002. Dunnett’s multiple comparisons test, saline vs FEN p = 0.3741, saline vs MDMA 7.5 p = 0.267, saline vs MDMA 15 **p = 0.0022, saline vs METH 2 ***p = 0.0003. **g.** Configuration of fiber photometry recording with GRAB DA in the NAc medial shell of wild-type mice. Scale = 100 um. **h.** Time course of bulk DA release triggered by each drug. **i.** Summary of area under the curve of DA release for each drug at the specified dose. Repeated measures one-way ANOVA, F_4,20_ = 11.06, ****p < 0.0001. Tukey’s multiple comparisons test, saline vs FEN 10 p = 0.9734, saline vs MDMA 7.5 p = 0.0812, saline vs MDMA 15 ***p = 0.0004, saline vs METH 2 ***p = 0.0009. **j.** Configuration of fiber photometry recording with GRAB 5-HT in the NAc medial shell of wild-type mice. Scale = 100 um. **k.** Time course of bulk 5-HT release triggered by each drug. **l.** Summary of area under the curve of 5-HT release for each drug at the specified dose. Repeated measures one-way ANOVA, F_2,6_ = 103.4, ****p < 0.0001. Tukey’s multiple comparisons test, saline vs MDMA 7.5 ****p < 0.001, saline vs METH 2 ***p = 0.0007, MDMA 7.5 vs METH 2 ***p = 0.0003.

Our behavioral data suggest that MDMA administered at 7.5 mg/kg triggers 5-HT release to levels that are prosocial but does not evoke DA release to levels that are rewarding. To evaluate the latter part of this hypothesis, we quantified DA release across drug conditions with fiber photometry recordings of DA release in the NAc medial shell, a key reward center, with the DA sensor GRAB_DA_ (Fig. 1g). Systemic injections of FEN (10 mg/kg), MDMA (7.5 mg/kg), or METH (2 mg/kg) led to varying levels of bulk DA release such that FEN triggered minimal release, METH evoked the largest release, and MDMA (7.5 mg/kg) at levels in between (Fig. 1h,i). MDMA at 15 mg/kg, however, produced DA release comparable to METH (Fig. 1h,i), consistent with its reinforcing behavioral effects.

We also addressed the assumption that, in this experimental preparation, MDMA produces dramatically higher levels of 5-HT release compared to METH, which has substantially lower (but detectable) action at SERT. We prepared a cohort of wild-type mice with the 5-HT sensor GRAB_5-HT_ in the NAc (Fig. 1j) to assess the 5-HT release evoked by the prosocial dose of MDMA (7.5 mg/kg) and the reinforcing dose of METH (2 mg/kg). MDMA produced a dramatically larger increase in 5-HT release than METH (Fig. 1k,l), as predicted. Together these data show that the dissociable effects of MDMA and METH on sociability and reinforcement are associated with differential DA and 5-HT release in the NAc. However, these data do not establish a causal relationship between these events. To probe the interaction between 5-HT and DA release in further detail, we next established the mechanisms by which MDMA evokes DA release.

### MDMA-evoked DA release and reward are governed through interactions with DAT

MDMA has at least two described mechanisms by which it releases DA: one involves reverse transport at DAergic terminals via DAT [8, 19, 40, 41]; another involves activity-dependent DA release regulated by action potentials in DAergic cell bodies of the ventral tegmental area (VTA) [22, 40, 42]. The contribution of these respective mechanisms may vary by dose of MDMA [22], and therefore may have differing contributions to DA-linked behaviors. We chose *Th*-Cre driver lines, rather than *Dat*-Cre, to study MDMA’s effects on DA release: although *Th*-Cre has less specificity for DAergic neurons [43], *Dat*-Cre lines show marked reduction in one of our proteins of interest, DAT [44]. We first evaluated the extent to which VTA activity is modulated by systemic MDMA. We injected AAV-DIO-GCaMP8m into the VTA of *Th*-Cre mice and recorded Ca^2+^ activity in DA neurons after injections of the prosocial and rewarding doses of MDMA (Fig. 2a,b). We observed a small but statistically significant increase in Ca^2+^ transient event frequency in VTA DA neurons after the high dose of MDMA (15 mg/kg; Fig. 2b). The same experiment performed in *Vgat*-Cre mice revealed minimal and inconsistent changes in VTA GABA neuron activity (Fig. 2c). In contrast, conditional deletion of DAT following tamoxifen injections (TMX; 75 mg/kg, ip) in floxed DAT mice crossed with *Th*-Cre^ERT2^ mice (DAT cKO mice) (Fig. 2d,e) led to a substantial reduction in MDMA-evoked DA release measured with GRAB_DA_ in the NAc (Fig. 2f,g). Absolute quantification of DA with microdialysis in the NAc confirmed the requirement of DAT for MDMA-induced DA release (Fig. 2h,i). These data suggest that MDMA-evoked DA release in the NAc derives predominantly from interactions with DAT, although small changes in DA cell firing could account for residual DA release at the 15 mg/kg dose.

**Figure 2.**
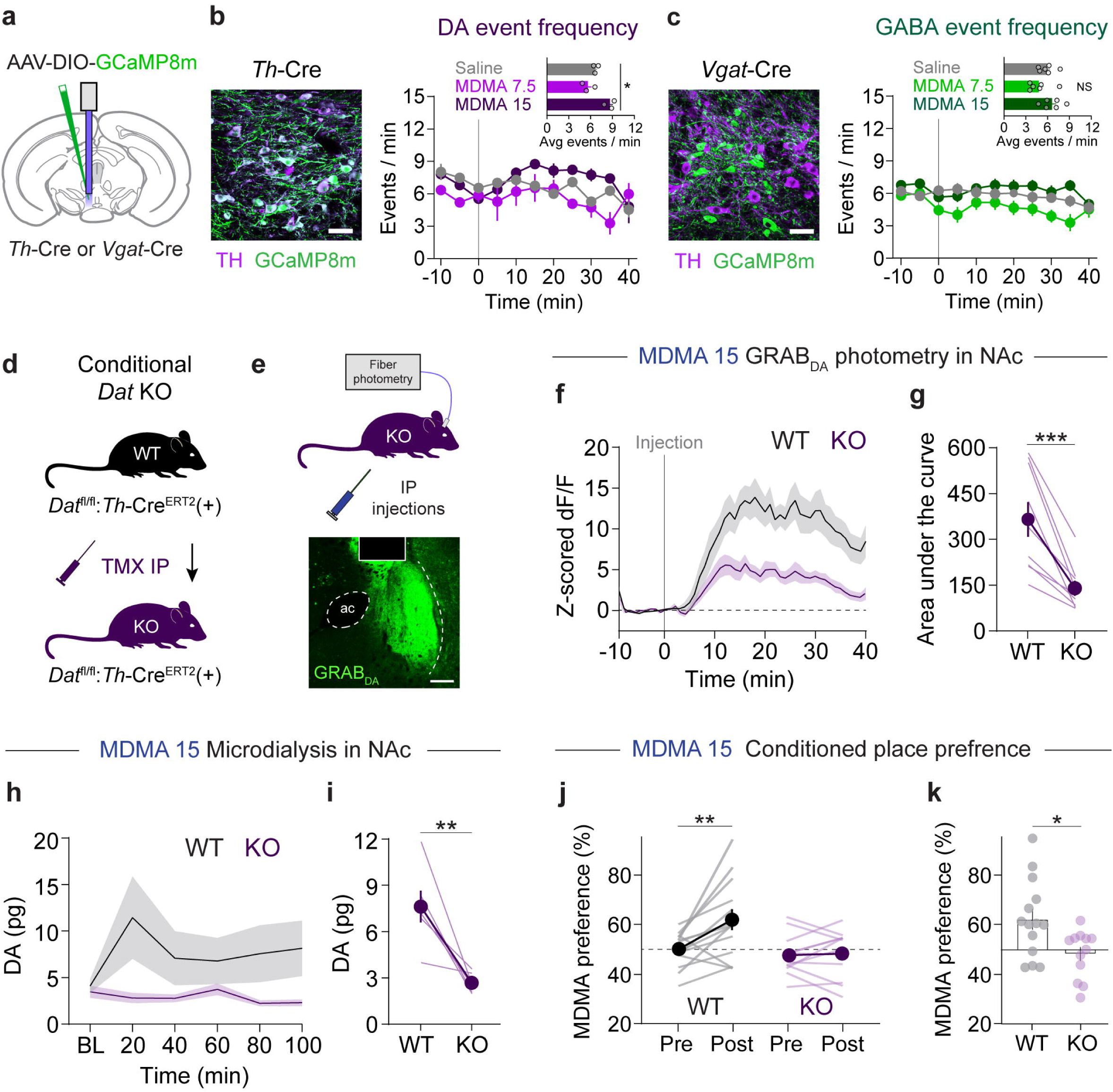
MDMA-evoked DA release and reward require DAT. **a.** Schematic of photometry recordings in the VTA of *Th*-Cre or *Vgat*-Cre mice. **b.** Left, image of VTA DA neurons infected with Cre-dependent GCaMP8m, scale = 20 um. Right, time course of DA neuron event frequency in 5 min bins. Inset, average frequency during the 15-25 min after MDMA injection. Repeated measures one-way ANOVA, F_2,2_ = 21.91, *p = 0.0341. Dunnett’s multiple comparisons test, saline vs MDMA 7.5 p = 0.3406, saline vs MDMA 15 p = 0.0752. **c**. Left, image of VTA GABA neurons infected with Cre-dependent GCaMP8m, scale = 20 um. Right, time course of GABA neuron event frequency in 5 min bins. Inset, average frequency during the 15-25 min after MDMA injection. Repeated measures one-way ANOVA, F_2,5_ = 4.965, p = 0.0615. Dunnett’s multiple comparisons test, saline vs MDMA 7.5 p = 0.0729, saline vs MDMA 15 p = 0.3354. **d.** Genetic strategy for generating conditional DAT knockout mice. *Dat*^fl/fl^:*Th*-Cre^ERT2^ mice were tested during a photometry recording and then injected with tamoxifen (TMX) to delete DAT. Mice were then tested again in a second photometry recording. **e.** Schematic of DA recording setup in knockout mice. **f.** Time course of DA release before (WT) and after (KO) deletion of DAT. **g.** Area under the curve of DA release in response to MDMA 15. Paired, two-tailed t-test, t_8_ = 5.137, ***p = 0.0009. **h.** Time course of DA release before (WT) and after (KO) deletion of DAT, as measured by microdialysis. **i.** DA concentrations after MDMA in both genotypes. Paired, two-tailed t-test, t_5_ = 4.205, **p = 0.0085. **j.** Effect of genetic deletion of DAT on MDMA (15 mg/kg) CPP. Two-way, repeated measures ANOVA, time x genotype interaction F_1,24_ = 6.506, *p = 0.0175. Sidak’s multiple comparisons test, Pre vs Post: WT **p = 0.0011, KO p = 0.9757, WT vs KO: Pre p = 0.757, Post **p = 0.0037. **k.** Summary and direct comparison of CPP performance. Unpaired, two-tailed t-test, t_24_ = 2.673, *p = 0.0133.

If DA release from MDMA is controlled through interactions with DAT, then deletion of DAT should also modify its reinforcing properties. We trained new groups of DAT cKO mice in a CPP procedure with the reinforcing dose of MDMA (15 mg/kg, ip). Conditional knockout of DAT significantly disrupted MDMA CPP (Fig. 2j,k), suggesting that MDMA’s interactions with DAT are also critical to its reinforcing properties.

To confirm genetic deletion of DAT in these mice, we also tested the effects of cocaine, a drug with a well-established requirement of DAT for its effects. Cocaine (15 mg/kg, ip) triggered a large increase in DA release in the NAc in WT mice, but this was greatly attenuated in DAT cKO mice (Supplementary Fig. 1a,b). In addition, cocaine at the same dose failed to establish CPP in DAT cKO mice (Supplementary Fig. 1c,d). These data confirm that DAT was functionally deleted from TH neurons, and that cocaine and MDMA share this mechanism to produce reinforcement.

### 5-HT2C receptors in NAc modulate MDMA reward and DA release

Our previous work demonstrated that viral mediated deletion of SERT abolished MDMA’s prosocial effects [10]. To confirm these findings with a similar yet orthogonal manipulation, we crossed floxed SERT mice to *Nes*-Cre^ERT2^ mice to generate SERT cKO mice where gene deletion preferentially occurs in 5-HTergic neurons after TMX administration (Fig. 3a) [12, 45]. Compared with control mice, SERT cKO mice do not show MDMA enhancement of prosocial behavior in the 3-CT (Fig. 3b). Surprisingly, MDMA could generate a robust CPP effect in SERT cKO mice at a dose (5 mg/kg, ip) that was 3-fold lower than required for wild-type mice (Fig. 3c,d), consistent with less selective pharmacological and genetic SERT manipulations [46, 47]. These data support the hypothesis that MDMA-induced 5-HT release promotes sociability yet actively limits MDMA reward, possibly by regulating MDMA-evoked DA release.

**Figure 3.**
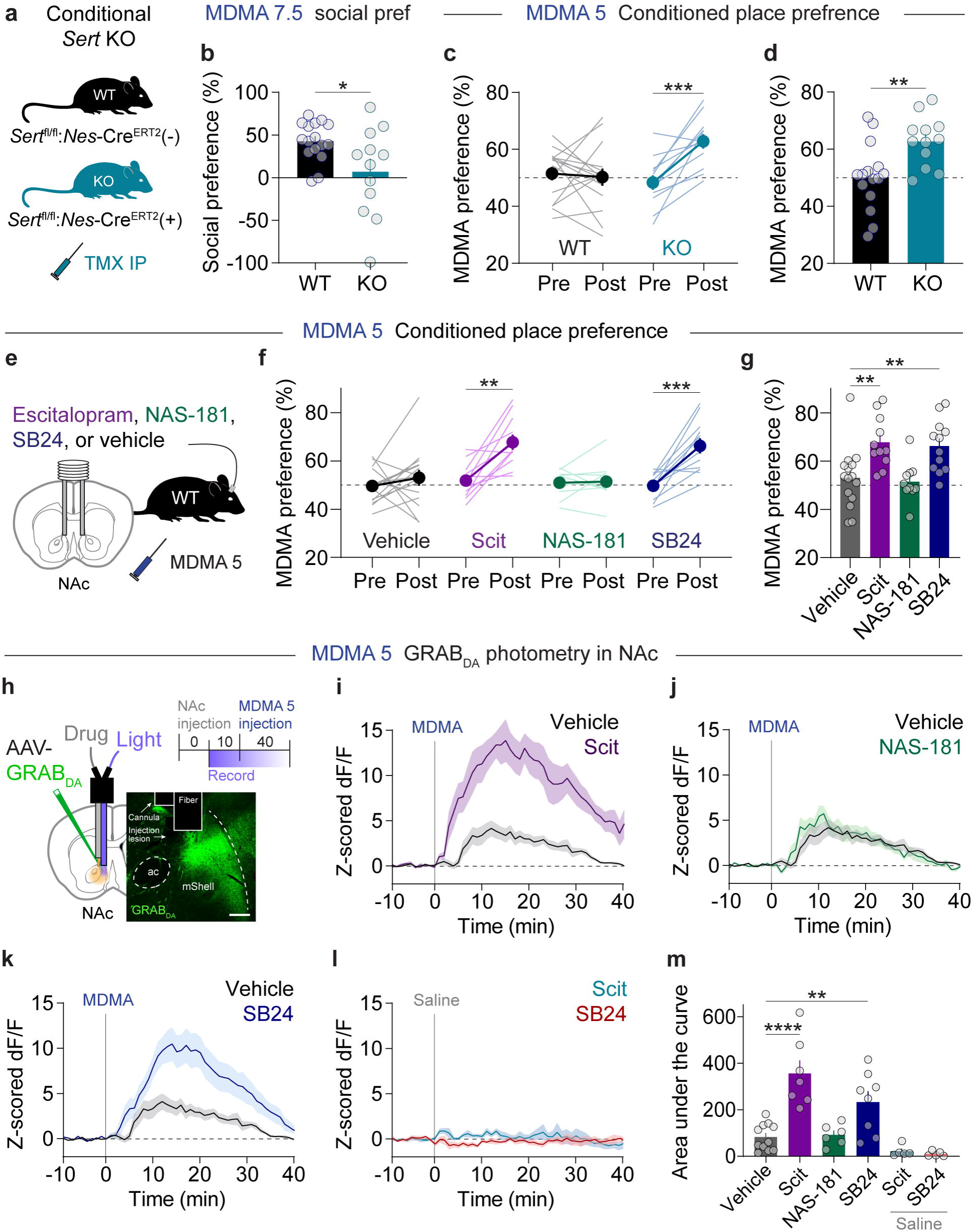
NAc 5-HT_2C_ receptors calibrate MDMA reward. **a.** Genetic strategy for conditional deletion of SERT. **b.** Effect of SERT deletion on 3-chamber social preference with MDMA (7.5 mg/kg). Unpaired, two-tailed t-test, t_26_ = 2.473, *p = 0.0203. **c.** SERT deletion enhanced CPP for MDMA with a subthreshold dose (5 mg/kg). Two-way, repeated measures ANOVA, time x genotype interaction F_1,25_ = 6.506, **p = 0.0027. Sidak’s multiple comparisons test, Pre vs Post: WT p = 0.9083, KO ***p = 0.0007, WT vs KO: Pre p = 0.6565, Post **p = 0.0032. **d.** Summary and direct comparison of CPP performance. Unpaired, two-tailed t-test, t_25_ = 3.098, **p = 0.0048. **e.** Experimental setup of microinfusions before subthreshold MDMA CPP. **f.** Effects of microinfusing each compound into the NAc on subthreshold MDMA CPP. Vehicle: paired, two-tailed t-test, t_14_ = 0.8939, p = 0.3865. Escitalopram (Scit): paired, two-tailed t-test, t_10_ = 4.385, **p = 0.0014. NAS-181: paired, two-tailed t-test, t_9_ = 0.2167, p = 0.8833. SB24: paired, two-tailed t-test, t_9_ = 5.220, ***p = 0.0003. **g.** Summary and direct comparison of CPP performance. Ordinary one-way ANOVA, F_3,44_ = 7.265, ***p = 0.0005. Dunnett’s multiple comparisons test, vehicle vs Scit **p = 0.0039, vehicle vs NAS-181 p = 0.9745, vehicle vs SB24 **p = 0.0084. **h.** Schematic of photometry recording setup for dual drug microinfusions into the NAc. Top right, experimental schedule. Scale = 100 um. **i.** Effect of intra-NAc infusion of Scit on bulk DA release evoked by MDMA (5 mg/kg). **j.** Effect of intra-NAc infusion of NAS-181 on bulk DA release evoked by MDMA (5 mg/kg). **k.** Effect of intra-NAc infusion of SB24 on bulk DA release evoked by MDMA (5 mg/kg). **l.** Effect of intra-NAc infusions of Scit and SB24 on bulk DA release after an injection of saline. **m.** Summary data of bulk DA levels after infusion of each compound. Ordinary one-way ANOVA, F_5,36_ = 14.54, ****p < 0.0001. Dunnett’s multiple comparisons test, vehicle vs Scit ****p < 0.0001, vehicle vs NAS-181 p = 0.9998, vehicle vs SB24 **p = 0.0049, vehicle vs Scit + saline p = 0.6553, vehicle vs SB24 + saline p = 0.4897.

5-HT neurons project widely across the brain [48, 49]. While we have observed SERT-dependent regulation of MDMA reward, it is unclear where in the brain this effect is mediated. To determine whether SERT engagement specifically in the NAc constrains MDMA’s reinforcing properties, we implanted bilateral guide cannulas targeting the NAc of wild-type mice and infused various 5-HT blockers during subthreshold MDMA CPP (Fig. 3e). We first infused the selective 5-HT reuptake inhibitor escitalopram (Scit, 1 μM) to prevent MDMA interactions with SERT [50]. This treatment significantly enhanced MDMA CPP (Fig. 3f,g), similar to SERT deletion. We previously found that 5-HT_1B_Rs in the NAc are required for the prosocial effects of 5-HT [12, 36, 51] and MDMA [10, 13], and therefore asked whether 5-HT released via MDMA also activates this receptor to modulate MDMA reinforcement. Surprisingly, infusion of the 5-HT_1B_R antagonist NAS-181 (1 μM) had no effect on subthreshold MDMA CPP (Fig. 3f,g). Previous reports have suggested that activation of 5-HT_2C_Rs can affect DA release in the NAc [52–54] and may account for the low abuse potential of classical psychedelics [26]. Therefore, we infused the 5-HT_2C_R antagonist SB242084 (SB24, 1 μM) into the NAc and observed significantly increased MDMA CPP, similar to Scit (Fig. 3f,g). These CPP data strongly suggest that MDMA engages 5-HT_2C_Rs that constrain its own rewarding effects.

If DA signaling in the NAc dictates MDMA’s rewarding properties and MDMA-evoked 5-HT release actively limits reward, then blocking MDMA’s ability to promote 5-HT signaling is predicted to also influence its effects on DA release. To determine whether 5-HT signaling in the NAc affects MDMA-evoked DA release, we implanted wild-type mice with a dual optical fiber-cannula device that allows for drug microinfusions directly into the photometry recording site (Fig. 3h). Mice were subjected to drug microinfusions through the cannula in their home cages and then immediately transferred to the photometry room for a DA recording during a systemic injection of MDMA (5 mg/kg, ip). We first infused Scit (1 μM) and detected substantial increases in bulk MDMA-evoked DA release compared with vehicle (Fig. 3i,m). As with CPP, infusion of the 5-HT_1B_R antagonist NAS-181 (1 μM) had no effect on MDMA-evoked DA release (Fig. 3j,m). By contrast, infusion of the 5-HT_2C_R antagonist SB24 (1 μM) significantly increased DA release after MDMA administration (Fig. 3k,m). Importantly, infusion of either Scit or SB24 after a systemic injection of saline had no effect on bulk DA levels (Fig. 3l,m). Altogether, these photometry recordings indicate that MDMA interactions at SERT in the NAc lead to 5-HT release, which activates 5-HT_2C_Rs that modulate DA release.

### (*R*)-MDMA possesses low addiction liability

Our data identify an interaction between MDMA-triggered DA and 5-HT in the NAc that calibrates both of its prosocial and reinforcing properties. Overcoming this 5-HT-mediated inhibition of reward and DA release required raising the dose of MDMA from 5mg/kg to 15mg/kg, the latter aligning with the effects of METH on CPP and DA release. Considering the need to screen candidate therapeutic compounds for abuse liability, our experiments so far show a rather predictable relationship between DA release and inducibility of CPP, suggesting that photometry-based DA measurement could be used to predict reinforcing characteristics of a novel entactogen. A potentially safer entactogen therapeutic would retain its 5-HT releasing properties but would release less DA over a range of doses.

The enantiomers of MDMA have different affinities for SERT and DAT [55–58]. Throughout this study, we have been administering the racemic (±*R*/*S*) mixture of MDMA enantiomers, an approximately 1:1 ratio of (*R*)-MDMA and S-MDMA. Of the two enantiomers, (*R*)-MDMA is a less potent monoamine releaser, however it more preferentially binds and releases neurotransmitter from SERT versus DAT [55–59], increases social contact in mice [39], and has lower abuse liability in a progressive ratio self-administration assay in rhesus monkeys [7]. Based on these properties, we predicted that administration of large doses of (*R*)-MDMA would act like (±)-MDMA at 7.5 mg/kg, i.e. is prosocial, not reinforcing, and triggers constrained levels of DA release. We examined the effects of (*R*)-MDMA on DA release in the NAc and observed a dose-dependent effect unlike (±)-MDMA. Compared with (±)-MDMA at 7.5 mg/kg, (*R*)-MDMA at 10 mg/kg evoked very low levels of DA release (Fig. 4a,b). (*R*)-MDMA at 20 mg/kg and 40 mg/kg were essentially identical to each other and (±)-MDMA at 7.5 mg/kg (Fig. 4a,b). These results suggest that (*R*)-MDMA (20 mg/kg) is not substantially reinforcing, similar to low dose (±)-MDMA (7.5 mg/kg). Consistent with this prediction, we could not detect any CPP for (*R*)-MDMA at 20 mg/kg (Fig. 4c). On the other hand, (*R*)-MDMA is predicted to retain its 5-HT releasing properties at a dose that induces prosocial behavior. We recorded 5-HT release in the NAc after an injection of (*R*)-MDMA (20 mg/kg, ip) and found statistically equivalent (and quantitatively greater) 5-HT release than that evoked by (±)-MDMA (7.5 mg/kg) (Fig. 4d,e). This increase in 5-HT release suggests that (*R*)-MDMA at 20 mg/kg, despite not being reinforcing, is highly prosocial. In the 3-CT, (*R*)-MDMA (20 mg/kg) evoked social preference that was significantly greater than saline and quantitatively greater than that induced by (±)-MDMA (7.5 mg/kg) (Fig. 4f and Fig. 1b,c). Altogether these data indicate that (*R*)-MDMA is prosocial but not reinforcing over a range of doses, thus likely possesses low potential for abuse.

**Figure 4.**
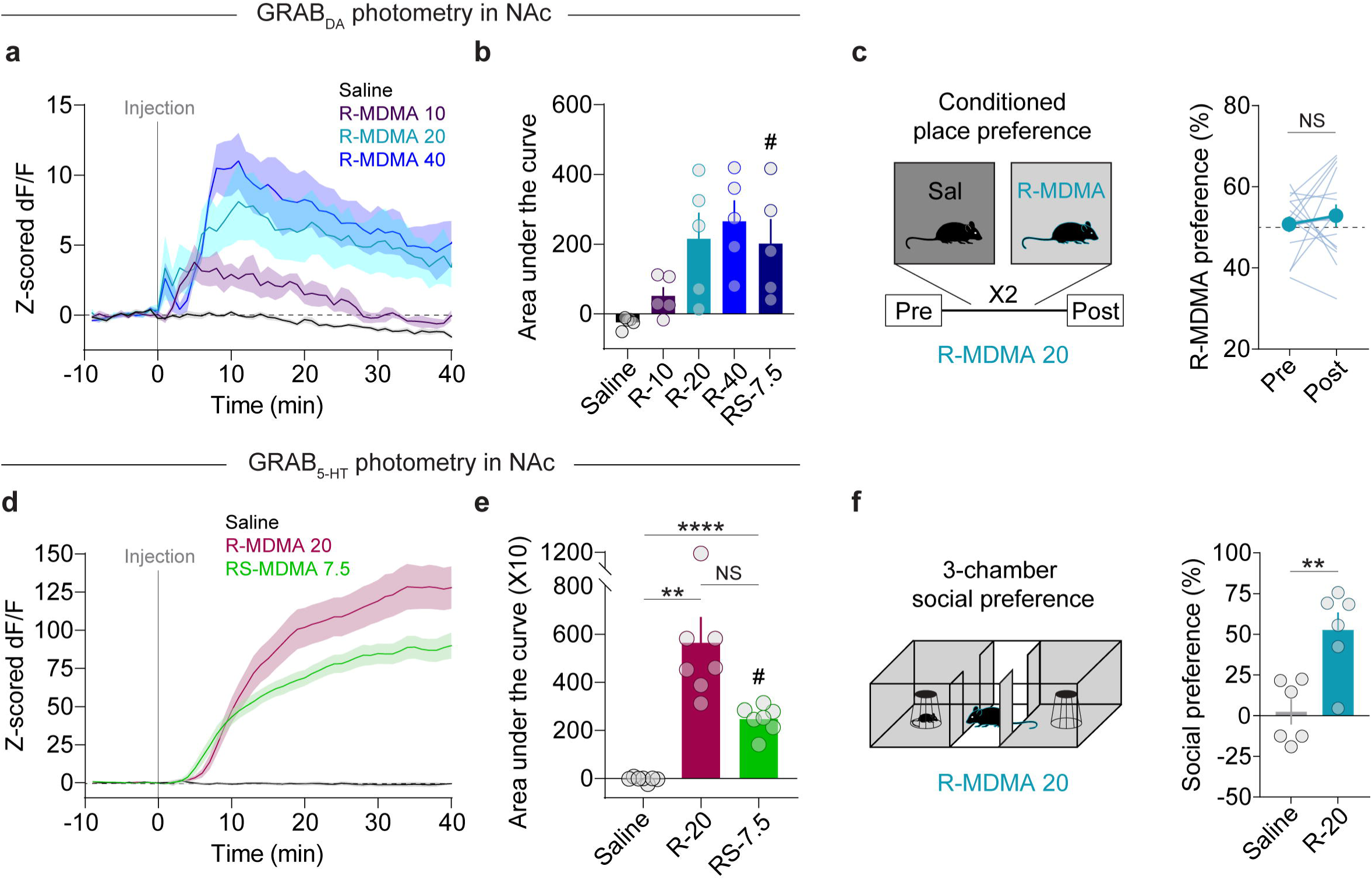
(*R*)-MDMA possesses desirable properties with less reinforcement. **a.** Time course of bulk DA release triggered by different doses of (*R*)-MDMA. **b.** Summary data of bulk DA levels after injection of each dose of (*R*)-MDMA, compared with racemic MDMA (7.5 mg/kg). Repeated measures one-way ANOVA, F_4,20_ = 14.54, **p = 0.0066. Dunnett’s multiple comparisons test, saline vs (*R*)-MDMA 10 p = 0.0535, saline vs (*R*)-MDMA 20 p = 0.0742, saline vs (*R*)-MDMA 40 *p = 0.0171, saline vs (±)-MDMA 7.5 p = 0.0777. ^#^(±)-MDMA 7.5 data from Figure 1. **c.** Left, schematic of CPP procedure. Right, magnitude of CPP produced with (*R*)-MDMA 20. Paired, two-tailed t-test, t_14_ = 0.6699, p = 0.5138. **d.** Time course of bulk 5-HT release triggered by (*R*)-MDMA (20 mg/kg). **e.** Summary data of bulk 5-HT levels after injection of (*R*)-MDMA (20 mg/kg), compared with racemic MDMA (7.5 mg/kg). Repeated measures one-way ANOVA, F_2,19_ = 17.17, **p = 0.0058. Tukey’s multiple comparisons test, saline vs (*R*)-MDMA 20 **p = 0.0048, saline vs (±)-MDMA 7.5 ****p < 0.0001, (*R*)-MDMA 20 vs (±)-MDMA 7.5 p = 0.1007. ^#^(±)-MDMA 7.5 data from Figure 1. **f.** Left, schematic of 3-chamber test with (*R*)-MDMA. Right, effect of (*R*)-MDMA (20 mg/kg) on social preference in the 3-chamber assay. Unpaired, two-tailed t-test, t_10_ = 3.777, **p = 0.0036.

## DISCUSSION

Developing novel entactogen-like drugs through preclinical assays requires a clear understanding of the mechanisms underlying MDMA’s presumed therapeutic effects, versus those that contribute to its misuse and abuse. We and others have previously found that MDMA’s prosocial effects in mice are well explained by release of 5-HT via SERT in the NAc, and activation of the 5-HT_1B_R. MDMA’s rewarding properties, independent of social context, are equally well explained by DA release in the NAc, consistent with the reward mechanism of other reinforcing drugs. In this study, we ask why MDMA appears to have a lower abuse liability than METH, whether that mechanism is one and the same as its prosocial mechanism, and whether preclinical assays based on this information can identify a novel potential therapeutic agent. We found that 5-HT release in the NAc does indeed account for the high dose threshold, relative to METH, for MDMA to induce CPP, a basic measure of drug reward. Though both the prosocial and reward-limiting effects of MDMA are linked to 5-HT release in the NAc, we found that they are mediated by separate 5-HT receptors. Unlike the 5-HT_1B_R-dependent prosocial effect of MDMA, the reward-limiting effect was only blocked by an intra-NAc infusion of a selective 5-HT_2C_R antagonist SB242084 [60]. During these experiments, we found a remarkably predictable relationship between NAc DA release, quantified by GRAB-DA fluorescence in the medial NAc, and the dose threshold required to elicit CPP. Using this information, we tested a range of doses of a putative 5-HT releasing entactogen, (*R*)-MDMA, finding that DA release appeared to plateau below levels that could be achieved with METH or high dose MDMA. Predictably, (*R*)-MDMA, produced prosocial effects but could not elicit CPP, suggesting it may be an entactogen with limited abuse liability.

In addition to a catalogue of effects on mood, anxiety, and appetitive drives, including social behavior, the 5-HTergic system has long been known to modulate the rewarding properties of reinforcing drugs. The flexibility of this control, reflected by the diversity of 5-HT receptors and brain regions implicated [21, 32, 61, 62], appears to match the variety of mechanisms engaged by various reinforcing drugs to release DA in the NAc, including disinhibition of GABAergic neurons of the VTA by morphine and ketamine [63, 64], direct activation of VTA neurons by ethanol and nicotine [63], DAT inhibition by cocaine, and reverse transport through DAT and VMAT by amphetamines [8]. Thus, a strategy to limit the abuse liability of phenethylamine / entactogen drugs like MDMA requires understanding the specific mechanism by which MDMA releases DA. Previous work points to both an activity-independent mechanism of DA release via DAT in the NAc [8, 40, 41] as well as an activity-dependent mechanism localized to the VTA [22, 40], although interpretation of prior work is limited by reliance on the constitutive deletion of DAT and use of nonselective modulators of activity, e.g. tetrodotoxin. Our data, which makes use of conditional DAT KO restricted to TH-expressing neurons and cell-type specific Ca^2+^ imaging in the VTA to index neural activity, indicates that DAT in the NAc accounts for the dominant proportion of DA release, and for MDMA drug reward.

While our data clearly localize 5-HTergic control over MDMA-evoked DA release to 5-HT_2C_R receptors expressed in the NAc, the cellular location of these 5-HT_2C_Rs is still unclear. Prior work has shown 5-HT_2C_R expression in the VTA that co-localizes with both DA and GABA neurons [65–67], and one prior study in rat found that a higher dose of the nonselective 5-HT_2B/2C_R antagonist SB206553 infused into VTA enhances NAc DA release evoked by MDMA[42]. Despite this complex expression pattern, our results with GCaMP8m recordings in the VTA suggest that this receptor is modulating DA release directly in the NAc. One possibility is that VTA GABA neurons express 5-HT_2C_Rs on their terminals in the NAc and these are engaged by 5-HT to release GABA onto local DA inputs. Despite coupling to Gq pathways and presumably increasing Ca^2+^ and excitability, it is possible 5-HT_2C_Rs receptors directly inhibit release in DA terminals through Ca^2+^-mediated potassium channel activation, atypical interactions with Gi biochemical pathways, or modulation of DAT function. Future work is necessary to determine the exact neural substrates 5-HT_2C_Rs receptors act on to modulate MDMA-evoked DA release.

The ease with which neurotransmitter release can now be quantified *in vivo* with fluorescent reporters like GRAB-DA and GRAB-5HT [68, 69] may reshape how preclinical drug discovery is performed, and has recently been applied to identifying ring-substituted cathinones with preferential 5-HT releasing properties [70]. While *in vitro* assays in expression and neuronal culture systems can measure the relative affinity amphetamine-derived drugs have for SERT or DAT [19, 70], our work showing the interplay between 5-HT and DA release via 5-HT_2C_R highlights the value of determining neurotransmitter release *in vivo*. In the course of identifying this interaction, we found a strong relationship between DA release in the NAc and the ability to induce a simple reward learning behavior, CPP, leading us to a simple test for relative DA-releasing efficacy of (*R*)-MDMA, an enantiomer that may have selective prosocial effects and less abuse liability in mice [71] and potentially humans [34]. By leveraging comparable methods in humans (e.g. radioligand-displacement PET imaging to measure DA and 5-HT release), a ‘fast-fail’ approach for clinical testing of novel candidate entactogens could be developed that incorporates biomarker-based proof-of-mechanism, as recently demonstrated in the evaluation of a novel kappa-opioid receptor antagonist for treatment of anhedonia [72].

### Limitations

The use of simple behavioral assays in mice, like 3-CT and CPP, are unlikely to represent the full range of human social behavior and patterns of drug misuse. Furthermore, photometric measurement of neurotransmitter release may be limited by a lack of specificity for one neurotransmitter, and an as-yet unclear relationship between the transients detected and neurotransmitter release kinetics. These caveats limit the interpretation of preclinical studies and form a strong argument for developing preclinical and clinical biomarker assays and simple behavioral readouts in parallel to maximize predictive power of these screening tools.

## ACKNOWLEDGEMENTS

We thank the entire Heifets, Eshel and Malenka Labs for helpful discussions, and the NIDA Drug Supply Program for supplying (±)-MDMA and (*R*)-MDMA. This work was supported by NIH grants K99 DA056573 (M.B.P.), K08 MH110610 (B.D.H.), R01 MH130591 (B.D.H.), P50 DA042012 (B.D.H. and R.C.M.), K08 MH123791 (N.E.), Brain & Behavior Research Foundation Young Investigator Award (N.E.), Burroughs Wellcome Fund Career Award for Medical Scientists (N.E.), Simons Foundation Bridge to Independence Award (N.E.), and a grant from the Stanford University Wu Tsai Neurosciences Institute (R.C.M.).

## SUPPLEMENTAL FIGURE LEGENDS

**Supplemental Figure 1. Analysis of DA release and cocaine CPP after DAT deletion.**

**a.** Time course of bulk DA release triggered by cocaine (15 mg/kg) in WT (pre TMX) and KO (post TMX) mice.

**b.** Quantification of area under the curve of DA release in response to cocaine.

**c.** CPP for cocaine (15 mg/kg) in WT and KO mice. Two-way, repeated measures ANOVA, time x genotype interaction F_1,13_ = 3.729, p = 0.0756. Sidak’s multiple comparisons test, Pre vs Post: WT *p = 0.0147, KO p = 0.9394.

**d.** Direct comparison of cocaine preference across genotypes. Unpaired, two-tailed t-test, t_13_ = 2.501, *p = 0.0265.

## REFERENCES

1. Mitchell JM, Bogenschutz M, Lilienstein A, Harrison C, Kleiman S, Parker-Guilbert K, et al. MDMA-assisted therapy for severe PTSD: a randomized, double-blind, placebo-controlled phase 3 study. Nat Med. 2021. 10 May 2021. 10.1038/s41591-021-01336-3.

2. Mitchell JM, Ot’alora G. M, van der Kolk B, Shannon S, Bogenschutz M, Gelfand Y, et al. MDMA-assisted therapy for moderate to severe PTSD: a randomized, placebo-controlled phase 3 trial. Nat Med. 2023:1–8.

3. Heifets BD, Olson DE. Therapeutic mechanisms of psychedelics and entactogens. Neuropsychopharmacology. 2024;49:104–118.

4. McCann UD, Ricaurte GA. Effects of MDMA on the Human Nervous System. The Effects of Drug Abuse on the Human Nervous System, Amsterdam: Elsevier; 2014. p. 475–497.

5. Shorter D, Hsieh J, Kosten TR. Pharmacologic management of comorbid post-traumatic stress disorder and addictions. Am J Addict. 2015;24:705–712.

6. De La Garza R, Fabrizio KR, Gupta A. Relevance of rodent models of intravenous MDMA self-administration to human MDMA consumption patterns. Psychopharmacology. 2007;189:425–434.

7. Wang Z, Woolverton WL. Estimating the relative reinforcing strength of (+/-)-3,4-methylenedioxymethamphetamine (MDMA) and its isomers in rhesus monkeys: comparison to (+)-methamphetamine. Psychopharmacology (Berl). 2007;189:483– 488.

8. Sulzer D. How addictive drugs disrupt presynaptic dopamine neurotransmission. Neuron. 2011;69:628–649.

9. Rothman RB, Baumann MH. Therapeutic and adverse actions of serotonin transporter substrates. Pharmacology & Therapeutics. 2002;95:73–88.

10. Heifets BD, Salgado JS, Taylor MD, Hoerbelt P, Cardozo Pinto DF, Steinberg EE, et al. Distinct neural mechanisms for the prosocial and rewarding properties of MDMA. Sci Transl Med. 2019;11:eaaw6435.

11. Nardou R, Lewis EM, Rothhaas R, Xu R, Yang A, Boyden E, et al. Oxytocin-dependent reopening of a social reward learning critical period with MDMA. Nature. 2019;569:116–120.

12. Walsh JJ, Christoffel DJ, Heifets BD, Ben-Dor GA, Selimbeyoglu A, Hung LW, et al. 5-HT release in nucleus accumbens rescues social deficits in mouse autism model. Nature. 2018;560:589–594.

13. Walsh JJ, Llorach P, Cardozo Pinto DF, Wenderski W, Christoffel DJ, Salgado JS, et al. Systemic enhancement of serotonin signaling reverses social deficits in multiple mouse models for ASD. Neuropsychopharmacology. 2021;46:2000–2010.

14. Rein B, Raymond K, Boustani C, Tuy S, Zhang J, St Laurent R, et al. MDMA enhances empathy-like behaviors in mice via 5-HT release in the nucleus accumbens. Sci Adv. 2024;10:eadl6554.

15. Kamilar-Britt P, Bedi G. The prosocial effects of 3,4-methylenedioxymethamphetamine (MDMA): Controlled studies in humans and laboratory animals. Neurosci Biobehav Rev. 2015;57:433–446.

16. Hysek CM, Schmid Y, Simmler LD, Domes G, Heinrichs M, Eisenegger C, et al. MDMA enhances emotional empathy and prosocial behavior. Soc Cogn Affect Neurosci. 2014;9:1645–1652.

17. Bershad AK, Miller MA, Baggott MJ, de Wit H. The effects of MDMA on socio-emotional processing: Does MDMA differ from other stimulants? J Psychopharmacol. 2016;30:1248–1258.

18. Vidal-Infer A, Roger-Sánchez C, Daza-Losada M, Aguilar MA, Miñarro J, Rodríguez-Arias M. Role of the dopaminergic system in the acquisition, expression and reinstatement of MDMA-induced conditioned place preference in adolescent mice. PLoS One. 2012;7:e43107.

19. Rothman RB, Baumann MH. Balance between dopamine and serotonin release modulates behavioral effects of amphetamine-type drugs. Ann N Y Acad Sci. 2006;1074:245–260.

20. Schenk S, Highgate Q. Methylenedioxymethamphetamine (MDMA): Serotonergic and dopaminergic mechanisms related to its use and misuse. J Neurochem. 2021;157:1714–1724.

21. Li Y, Simmler LD, Van Zessen R, Flakowski J, Wan J-X, Deng F, et al. Synaptic mechanism underlying serotonin modulation of transition to cocaine addiction. Science. 2021;373:1252–1256.

22. Federici M, Sebastianelli L, Natoli S, Bernardi G, Mercuri NB. Electrophysiologic Changes in Ventral Midbrain Dopaminergic Neurons Resulting from (+/−) -3,4-Methylenedioxymethamphetamine (MDMA—“Ecstasy”). Biological Psychiatry. 2007;62:680–686.

23. O’Dell LE, Parsons LH. Serotonin1B receptors in the ventral tegmental area modulate cocaine-induced increases in nucleus accumbens dopamine levels. J Pharmacol Exp Ther. 2004;311:711–719.

24. Fletcher PJ, Chintoh AF, Sinyard J, Higgins GA. Injection of the 5-HT2C receptor agonist Ro60-0175 into the ventral tegmental area reduces cocaine-induced locomotor activity and cocaine self-administration. Neuropsychopharmacology. 2004;29:308–318.

25. Navailles S, Moison D, Cunningham KA, Spampinato U. Differential regulation of the mesoaccumbens dopamine circuit by serotonin2C receptors in the ventral tegmental area and the nucleus accumbens: an in vivo microdialysis study with cocaine. Neuropsychopharmacology. 2008;33:237–246.

26. Canal CE, Murnane KS. The serotonin 5-HT2C receptor and the non-addictive nature of classic hallucinogens. J Psychopharmacol. 2017;31:127–143.

27. Fletcher PJ, Azampanah A, Korth KM. Activation of 5-HT(1B) receptors in the nucleus accumbens reduces self-administration of amphetamine on a progressive ratio schedule. Pharmacol Biochem Behav. 2002;71:717–725.

28. Fletcher PJ, Korth KM. Activation of 5-HT1B receptors in the nucleus accumbens reduces amphetamine-induced enhancement of responding for conditioned reward. Psychopharmacology (Berl). 1999;142:165–174.

29. Zayara AE, McIver G, Valdivia PN, Lominac KD, McCreary AC, Szumlinski KK. Blockade of nucleus accumbens 5-HT2A and 5-HT2C receptors prevents the expression of cocaine-induced behavioral and neurochemical sensitization in rats. Psychopharmacology (Berl). 2011;213:321–335.

30. Auclair AL, Cathala A, Sarrazin F, Depoortère R, Piazza PV, Newman-Tancredi A, et al. The central serotonin2B receptor: a new pharmacological target to modulate the mesoaccumbens dopaminergic pathway activity. J Neurochem. 2010;114:1323–1332.

31. Doly S, Valjent E, Setola V, Callebert J, Hervé D, Launay J-M, et al. Serotonin 5-HT2B receptors are required for 3,4-methylenedioxymethamphetamine-induced hyperlocomotion and 5-HT release in vivo and in vitro. J Neurosci. 2008;28:2933– 2940.

32. Howell LL, Cunningham KA. Serotonin 5-HT2 receptor interactions with dopamine function: implications for therapeutics in cocaine use disorder. Pharmacol Rev. 2015;67:176–197.

33. Devroye C, Filip M, Przegaliński E, McCreary AC, Spampinato U. Serotonin2C receptors and drug addiction: focus on cocaine. Exp Brain Res. 2013;230:537–545.

34. Straumann I, Avedisian I, Klaiber A, Varghese N, Eckert A, Rudin D, et al. Acute effects of R-MDMA, S-MDMA, and racemic MDMA in a randomized double-blind cross-over trial in healthy participants. Neuropsychopharmacology. 2024. 23 August 2024. 10.1038/s41386-024-01972-6.

35. Chen X, Ye R, Gargus JJ, Blakely RD, Dobrenis K, Sze JY. Disruption of Transient Serotonin Accumulation by Non-Serotonin-Producing Neurons Impairs Cortical Map Development. Cell Rep. 2015;10:346–358.

36. Wu X, Morishita W, Beier KT, Heifets BD, Malenka RC. 5-HT modulation of a medial septal circuit tunes social memory stability. Nature. 2021;599:96–101.

37. Pomrenze MB, Cardozo Pinto DF, Neumann PA, Llorach P, Tucciarone JM, Morishita W, et al. Modulation of 5-HT release by dynorphin mediates social deficits during opioid withdrawal. Neuron. 2022;110:4125–4143.e6.

38. Esaki H, Sasaki Y, Nishitani N, Kamada H, Mukai S, Ohshima Y, et al. Role of 5-HT1A receptors in the basolateral amygdala on 3,4-methylenedioxymethamphetamine-induced prosocial effects in mice. Eur J Pharmacol. 2023;946:175653.

39. Curry DW, Young MB, Tran AN, Daoud GE, Howell LL. Separating the agony from ecstasy: R(-)-3,4-methylenedioxymethamphetamine has prosocial and therapeutic-like effects without signs of neurotoxicity in mice. Neuropharmacology. 2018;128:196–206.

40. Gudelsky GA, Nash JF. Carrier-mediated release of serotonin by 3,4-methylenedioxymethamphetamine: implications for serotonin-dopamine interactions. J Neurochem. 1996;66:243–249.

41. Hagino Y, Takamatsu Y, Yamamoto H, Iwamura T, L. Murphy D, R. Uhl G, et al. Effects of MDMA on Extracellular Dopamine and Serotonin Levels in Mice Lacking Dopamine and/or Serotonin Transporters. CN. 2011;9:91–95.

42. Bankson MG, Yamamoto BK. Serotonin-GABA interactions modulate MDMA-induced mesolimbic dopamine release. J Neurochem. 2004;91.

43. Lammel S, Steinberg EE, Földy C, Wall NR, Beier K, Luo L, et al. Diversity of transgenic mouse models for selective targeting of midbrain dopamine neurons. Neuron. 2015;85:429–438.

44. Bäckman CM, Malik N, Zhang Y, Shan L, Grinberg A, Hoffer BJ, et al. Characterization of a mouse strain expressing Cre recombinase from the 3’ untranslated region of the dopamine transporter locus. Genesis. 2006;44:383–390.

45. Sun M-Y, Yetman MJ, Lee T-C, Chen Y, Jankowsky JL. Specificity and efficiency of reporter expression in adult neural progenitors vary substantially among nestin-CreER(T2) lines. J Comp Neurol. 2014;522:1191–1208.

46. Oakly AC, Brox BW, Schenk S, Ellenbroek BA. A genetic deletion of the serotonin transporter greatly enhances the reinforcing properties of MDMA in rats. Mol Psychiatry. 2014;19:534–535.

47. Roger-Sánchez C, Aguilar MA, Manzanedo C, Miñarro J, Rodríguez-Arias M. Neurochemical substrates of MDMA reward: effects of the inhibition of serotonin reuptake on the acquisition and reinstatement of MDMA-induced CPP. Curr Pharm Des. 2013;19:7050–7064.

48. Ren J, Friedmann D, Xiong J, Liu CD, Ferguson BR, Weerakkody T, et al. Anatomically Defined and Functionally Distinct Dorsal Raphe Serotonin Sub-systems. Cell. 2018;175:472–487.e20.

49. Cardozo Pinto DF, Yang H, Pollak Dorocic I, de Jong JW, Han VJ, Peck JR, et al. Characterization of transgenic mouse models targeting neuromodulatory systems reveals organizational principles of the dorsal raphe. Nat Commun. 2019;10:4633.

50. Gudelsky GA, Yamamoto BK. Actions of 3,4-methylenedioxymethamphetamine (MDMA) on cerebral dopaminergic, serotonergic and cholinergic neurons. Pharmacology Biochemistry and Behavior. 2008;90:198–207.

51. Dölen G, Darvishzadeh A, Huang KW, Malenka RC. Social reward requires coordinated activity of nucleus accumbens oxytocin and serotonin. Nature. 2013;501:179–184.

52. Browne CJ, Ji X, Higgins GA, Fletcher PJ, Harvey-Lewis C. Pharmacological Modulation of 5-HT2C Receptor Activity Produces Bidirectional Changes in Locomotor Activity, Responding for a Conditioned Reinforcer, and Mesolimbic DA Release in C57BL/6 Mice. Neuropsychopharmacol. 2017;42:2178–2187.

53. Navailles S, De Deurwaerdère P, Porras G, Spampinato U. In Vivo Evidence that 5-HT2C Receptor Antagonist but not Agonist Modulates Cocaine-Induced Dopamine Outflow in the Rat Nucleus Accumbens and Striatum. Neuropsychopharmacol. 2004;29:319–326.

54. Zhang G, Wu X, Zhang Y-M, Liu H, Jiang Q, Pang G, et al. Activation of serotonin 5-HT2C receptor suppresses behavioral sensitization and naloxone-precipitated withdrawal symptoms in morphine-dependent mice. Neuropharmacology. 2016;101:246–254.

55. Steele TD, Nichols DE, Yim GK. Stereochemical effects of 3,4-methylenedioxymethamphetamine (MDMA) and related amphetamine derivatives on inhibition of uptake of [3H]monoamines into synaptosomes from different regions of rat brain. Biochem Pharmacol. 1987;36:2297–2303.

56. Setola V, Hufeisen SJ, Grande-Allen KJ, Vesely I, Glennon RA, Blough B, et al. 3,4-Methylenedioxymethamphetamine (MDMA, “Ecstasy”) Induces Fenfluramine-Like Proliferative Actions on Human Cardiac Valvular Interstitial Cells in Vitro. Mol Pharmacol. 2003;63:1223–1229.

57. Verrico CD, Miller GM, Madras BK. MDMA (Ecstasy) and human dopamine, norepinephrine, and serotonin transporters: implications for MDMA-induced neurotoxicity and treatment. Psychopharmacology (Berl). 2007;189:489–503.

58. Huot P, Johnston TH, Lewis KD, Koprich JB, Reyes MG, Fox SH, et al. Characterization of 3,4-Methylenedioxymethamphetamine (MDMA) Enantiomers *In Vitro* and in the MPTP-Lesioned Primate: *R* -MDMA Reduces Severity of Dyskinesia, Whereas *S* -MDMA Extends Duration of ON-Time. J Neurosci. 2011;31:7190–7198.

59. Hiramatsu M, Cho AK. Enantiomeric differences in the effects of 3,4-methylenedioxymethamphetamine on extracellular monoamines and metabolites in the striatum of freely-moving rats: an in vivo microdialysis study. Neuropharmacology. 1990;29.

60. Kennett GA, Wood MD, Bright F, Trail B, Riley G, Holland V, et al. SB 242084, a Selective and Brain Penetrant 5-HT2C Receptor Antagonist. Neuropharmacology. 1997;36:609–620.

61. Porras G, Di Matteo V, Fracasso C, Lucas G, De Deurwaerdère P, Caccia S, et al. 5-HT2A and 5-HT2C/2B Receptor Subtypes Modulate Dopamine Release Induced in Vivo by Amphetamine and Morphine in Both the Rat Nucleus Accumbens and Striatum. Neuropsychopharmacol. 2002;26:311–324.

62. Porrino LJ, Ritz MC, Goodman NL, Sharpe LG, Kuhar MJ, Goldberg SR. Differential effects of the pharmacological manipulation of serotonin systems on cocaine and amphetamine self-administration in rats. Life Sciences. 1989;45:1529– 1535.

63. Juarez B, Han M-H. Diversity of Dopaminergic Neural Circuits in Response to Drug Exposure. Neuropsychopharmacology. 2016;41:2424–2446.

64. Simmler LD, Li Y, Hadjas LC, Hiver A, van Zessen R, Lüscher C. Dual action of ketamine confines addiction liability. Nature. 2022;608:368–373.

65. Bubar MJ, Stutz SJ, Cunningham KA. 5-HT2C Receptors Localize to Dopamine and GABA Neurons in the Rat Mesoaccumbens Pathway. PLOS ONE. 2011;6:e20508.

66. Bubar MJ, Cunningham KA. Distribution of serotonin 5-HT2C receptors in the ventral tegmental area. Neuroscience. 2007;146:286–297.

67. Eberle-Wang K, Mikeladze Z, Uryu K, Chesselet M-F. Pattern of expression of the serotonin2C receptor messenger RNA in the basal ganglia of adult rats. J Comp Neurol. 1997;384:233–247.

68. Sun F, Zhou J, Dai B, Qian T, Zeng J, Li X, et al. Next-generation GRAB sensors for monitoring dopaminergic activity in vivo. Nat Methods. 2020;17:1156–1166.

69. Wan J, Peng W, Li X, Qian T, Song K, Zeng J, et al. A genetically encoded sensor for measuring serotonin dynamics. Nat Neurosci. 2021;24:746–752.

70. Mayer FP, Niello M, Cintulova D, Sideromenos S, Maier J, Li Y, et al. Serotonin-releasing agents with reduced off-target effects. Mol Psychiatry. 2023;28:722–732.

71. Pitts EG, Curry DW, Hampshire KN, Young MB, Howell LL. (±)-MDMA and its enantiomers: potential therapeutic advantages of R(-)-MDMA. Psychopharmacology (Berl). 2018;235:377–392.

72. Krystal AD, Pizzagalli DA, Smoski M, Mathew SJ, Nurnberger J, Lisanby SH, et al. A randomized proof-of-mechanism trial applying the ‘fast-fail’ approach to evaluating κ-opioid antagonism as a treatment for anhedonia. Nat Med. 2020;26:760–768.

