## Supplementary figures and images for "5-HT_2C_ receptors in the nucleus accumbens constrain the rewarding effects of MDMA"

### Supplemental Figure 1

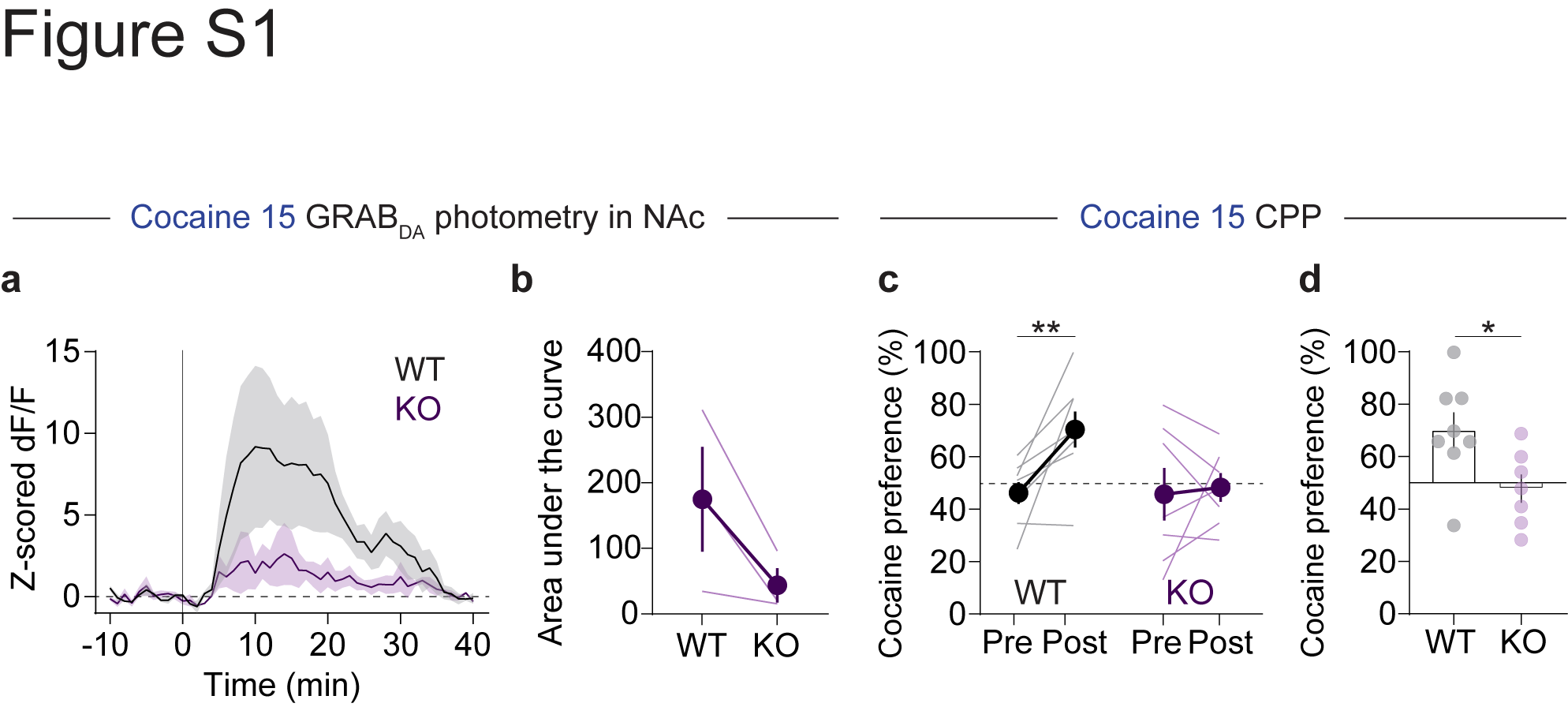
